# Evidence for natural hybridisation and novel *Wolbachia* strain superinfections in the *Anopheles gambiae* complex from Guinea

**DOI:** 10.1101/772855

**Authors:** Claire L Jeffries, Cintia Cansado-Utrilla, Abdoul H Beavogui, Caleb Stica, Eugene K Lama, Mojca Kristan, Seth R Irish, Thomas Walker

## Abstract

*Wolbachia*, a widespread bacterium which can influence mosquito-borne pathogen transmission, has recently been detected within *Anopheles (An.)* species that are malaria vectors in Sub-Saharan Africa. Although studies have reported *Wolbachia* strains in the *An. gambiae* complex, apparent low density and prevalence rates require confirmation. In this study, wild *Anopheles* mosquitoes collected from two regions of Guinea were investigated. In contrast to previous studies, RNA was extracted from adult females (n=516) to increase the chances for detection of actively expressed *Wolbachia* genes, determine *Wolbachia* prevalence rates and estimate relative strain densities. Molecular confirmation of mosquito species and *Wolbachia* Multilocus sequence typing (MLST) were carried out to analyse phylogenetic relationships of mosquito hosts and newly discovered *Wolbachia* strains. Strains were detected in *An. gambiae* s.s. (prevalence rates of 0.0-2.8%) from the Faranah region, *An. melas* (prevalence rate of 11.6% - 16/138) and hybrids between these two species (prevalence rate of 40.0% - 6/15) from Senguelen in the Maferinyah region. Furthermore, a novel high-density strain, termed *w*AnsX, was found in an unclassified *Anopheles* species. The discovery of novel *Wolbachia* strains (particularly in members, and hybrids, of the *An. gambiae* complex) provides further candidate strains that could be used for future *Wolbachia*-based malaria biocontrol strategies.

## 2. Introduction

*Wolbachia* endosymbiotic bacteria are estimated to infect ~40% of insect species [1] and natural infections have been shown to have inhibitory effects on human arboviruses in mosquitoes [2–4]. High density *Wolbachia* strains have been utilised for mosquito biocontrol strategies targeting arboviruses as they induce synergistic phenotypic effects. *Wolbachia* strains that have been transinfected into *Aedes (Ae.) aegypti* and *Ae. albopictus* induce inhibitory effects on arboviruses, with maternal transmission and cytoplasmic incompatibility (CI) enabling introduced strains to spread through populations [5–13]. The successful release and establishment of *Wolbachia*-transinfected *Ae. aegypti* populations in Cairns, Australia [14] was followed by further evidence of strong inhibitory effects on arboviruses from field populations [15]. Further studies in Australia [16, 17] and Kuala Lumpur, Malaysia [18] have now shown that *Wolbachia* frequencies have remained stable since initial releases and there is a reduction in human dengue incidence (case notifications) in the release sites.

The potential for *Wolbachia* to be used for biocontrol strategies targeting malaria transmission by *Anopheles* species has also been postulated [19] and initial laboratory experiments demonstrated that transient infections in *An. gambiae* reduce the density of *Plasmodium (P.) falciparum* parasites [20]. However, as with arboviruses there is variability in the level of inhibition of malaria parasites for different *Wolbachia* strains in different mosquito species [21–23]. A major step forward was achieved through the transinfection of a *Wolbachia* strain from *Ae. albopictus* (wAlbB) into An. stephensi and the confirmation of P. falciparum inhibition [24]. The interest in using *Wolbachia* for biocontrol strategies targeting malaria transmission in *Anopheles* mosquitoes has further increased due to the detection of natural strains of *Wolbachia* residing in numerous malaria vectors of Sub-Saharan Africa [25–29]. The *An. gambiae* complex, which consists of multiple morphologically indistinguishable species including several major malaria vector species, appears to contain diverse *Wolbachia* strains (collectively named *w*Anga) at both low prevalence and low infection densities [25, 26, 28–31]. In contrast, the recently discovered *w*AnM and *w*AnsA strains, found in *An. moucheti* and *An.* species A respectively, are higher density infections that dominate the mosquito microbiome [26].

Interestingly, the presence of *Wolbachia* strains in *Anopheles* was inversely correlated to other bacteria species such as *Asaia* that are stably associated with several species [32–34]. Evidence for this ‘mutual exclusion’ between bacterial species in *Anopheles* was also present from analysis of field collected mosquitoes from multiple countries in Sub-Saharan Africa [26]. In this study, we collected wild *Anopheles* mosquitoes from two regions of Guinea in June-July 2018 and characterised the natural *Wolbachia* strains to provide further evidence for the presence of these endosymbionts in malaria vectors. In contrast to previous studies, we extracted RNA to make any detection of *Wolbachia* more likely to be from actively expressed *Wolbachia* genes and undertook qRT-PCR analysis to compare *Wolbachia* densities. Phylogenetic analysis revealed the presence of novel strains in *An. melas*, *An. gambiae* s.s.*-melas* hybrids (including *Wolbachia* superinfections within individual mosquitoes) and an unclassified *Anopheles* species.

## 3. Materials and Methods

### Study sites & collection methods

*Anopheles* adult mosquitoes were collected in 2018 from two regions (sub-prefectures) in Guinea; Faranah and Maferinyah. Human landing catches (HLCs) and larval dipping were conducted in three villages in the Faranah Prefecture; Balayani (10.1325, −10.7443), Foulaya (10.144633, - 10.749717), and Tindo (9.9612230, −10.7016560) [35]. Three districts were selected for mosquito collections in the Maferinyah sub-prefecture using a variety of traps [36]. BG sentinel 2 traps (BG2) (Biogents), CDC light traps (John W. Hock), gravid traps (BioQuip) and stealth traps (John W. Hock) were used to sample adult mosquitoes in Maferinyah Centre I (09.54650, −013.28160), Senguelen (09.41150, −013.37564) and Fandie (09.53047, −013.24000). Mosquitoes collected from traps and HLCs were morphologically identified using keys and stored in RNAlater^®^ (Invitrogen) at −70°C [35, 36].

### RNA extraction and generation of cDNA

RNA was extracted from individual whole female mosquitoes using Qiagen 96 RNeasy Kits according to manufacturer’s instructions and a Qiagen Tissue Lyser II (Hilden, Germany) with a 5mm stainless steel bead (Qiagen) to homogenise mosquitoes. RNA was eluted in 45 μL of RNase-free water and stored at −70°C. RNA was reverse transcribed into complementary DNA (cDNA) using an Applied Biosystems High Capacity cDNA Reverse Transcription kit. A final volume of 20 μL contained 10 μL RNA, 2 μL 10X RT buffer, 0.8 μL 25X dNTP (100 mM), 2 μL 10X random primers, 1μL reverse transcriptase and 4.2 μL nuclease-free water. Reverse transcription was undertaken in a Bio-Rad T100 Thermal Cycler as follows: 25°C for 10min, 37°C for 120min and 85°C for 5min and cDNA stored at –20°C.

### Molecular mosquito species identification

Species identification of the *An. gambiae* complex was initially undertaken using diagnostic species-specific PCR assays targeting the ribosomal intergenic spacer (IGS)[37] and SINE200 insertion[38] to distinguish between the morphologically indistinguishable sibling species. To confirm species identification for samples of interest and samples that could not be identified by species-specific PCR, Sanger sequencing and phylogenetic analysis was performed for PCR products from a range of gene targets including ribosomal IGS and internal transcribed spacer 2 (ITS2)[39] and mitochondrial cytochrome c oxidase subunit 1 (*COI*)[40], cytochrome c oxidase subunit 2 (*COII*)[41] and NADH dehydrogenase subunits 4 and 5 (*ND4-ND5*)[42]. Where ITS2 PCR products for a particular sample were not successfully generated, or the sequencing generated was not of sufficient quality for onward analysis, a slight modification to the ITS2 primers was used to attempt to increase the success of amplification and sequencing. Alternative ITS2 primers adjusted from those published[39] were ITS2A-CJ: 5’-TGTGAACTTGCAGGACACAT-3’ and ITS2B-CJ: 5’-TATGCTTAAATTYAGGGGGT-3’. For confirmation of *Culex (Cx.) watti* - a species collected in the same location and used for comparative *Wolbachia* density analysis - a different fragment of the mitochondrial cytochrome c oxidase subunit 1 (*COI*) gene [43] was sequenced given the lack of available sequences in certain regions for this species and to optimise sequencing quality and species discrimination. PCR reactions for IGS, SINE200, ITS2 and COI were prepared as previously described[36]. For COII amplification, PCR reactions were prepared using 10 μL of Phire Hot Start II PCR Master Mix (Thermo Scientific™) with a final concentration of 1 μM of each primer, 1 μL of PCR grade water and 2 μL template cDNA, to a final reaction volume of 20 μL. PCR reactions were carried out in a Bio-Rad T100 Thermal Cycler and cycling was 98°C for 30 sec followed by 34 cycles of 98°C for 5 sec, 55°C for 5 sec, 72°C for 30 sec followed by 72°C for 1 min. For *ND4-ND5* PCR reactions were prepared using 10 μL of HotStart Taq 2x Master Mix (New England BioLabs^®^) with a final concentration of 2 μM of each primer, 1 μL of PCR grade water and 2 μL template cDNA, to a final reaction volume of 20 μL. PCR reactions were carried out in a Bio-Rad T100 Thermal Cycler and cycling was 95°C for 30 sec followed by 35 cycles of 95°C for 30 sec, 53°C for 60 sec, 68°C for 90 sec followed by 68°C for 5 min. PCR products were separated and visualised using 2% E-Gel EX agarose gels (Invitrogen) with SYBR safe and an Invitrogen E-Gel iBase Real-Time Transilluminator.

### *Wolbachia* detection and amplification of *Wolbachia* genes

*Wolbachia* detection was first undertaken targeting the conserved *Wolbachia* genes previously shown to amplify a wide diversity of strains; *16S rRNA* gene using primers W-Spec-16S-F: 5’-CATACCTATTCGAAGGGATA-3’ and W-Spec-16s-R: 5’-AGCTTCGAGTGAAACCAATTC-3 [44] and *Wolbachia* surface protein (*wsp*) gene using primers *wsp*81F: 5’-TGGTCCAATAAGTGATGAAGAAAC-3’ and *wsp*691R: 5’-AAAAATTAAACGCTACTCCA-3’ [45]. PCR analysis was also undertaken on cDNA to determine if there was any evidence for the presence of CI-inducing genes *CifA* (primers 5’-TGTGGTAGGGAAGGAAAGAGGAAA-3’, 5’-ATTCCAAGGACCATCACCTACAGA-3’) and *CifB* (primers 5’-TGCGAGAGATTAGAGGGCAAAATC-3’, 5’-CCTAAGAAGGCTAATCTCAGACGC-3’) [46]. Multilocus strain typing (MLST) was undertaken to characterize *Wolbachia* strains using the sequences of five conserved genes as molecular markers to genotype each strain. In brief, 450-500 base pair fragments of the *gatB, coxA, hcpA, ftsZ* and *fbpA Wolbachia* genes were amplified from individual *Wolbachia*-infected mosquitoes using previously optimised protocols [47, 48]. Primers used were as follows: gatB_F1: 5’-GAKTTAAAYCGYGCAGGBGTT-3’, gatB_R1: 5’-TGGYAAYTCRGGYAAAGATGA-3’, coxA_F1: 5’-TTGGRGCRATYAACTTTATAG-3’, coxA_R1: 5’-CTAAAGACTTTKACRCCAGT-3’, hcpA_F1: 5’-GAAATARCAGTTGCTGCAAA-3’, hcpA_R1: 5’-GAAAGTYRAGCAAGYTCTG-3’, ftsZ_F1: 5’-ATYATGGARCATATAAARGATAG-3’, ftsZ_R1: 5’-TCRAGYAATGGATTRGATAT-3’, fbpA_F1: 5’-GCTGCTCCRCTTGGYWTGAT-3’ and fbpA_R1: 5’-CCRCCAGARAAAAYYACTATTC-3’ with the addition of M13 adaptors. If no amplification was detected using standard primers, further PCR analysis was undertaken using degenerate primer sets, with or without M13 adaptors [47]. In selected *An. melas* specimens where *Wolbachia *16S rRNA** Sanger sequencing (detailed below) indicated the possibility of superinfections, further MLST testing was carried out utilising *Wolbachia* Supergroup A and B strain specific primers [47]. PCR reactions were prepared using 10 μL of Phire Hot Start II PCR Master Mix (Thermo Scientific™) with a final concentration of 1 μM of each primer, 1 μL of PCR grade water and 2 μL template cDNA, to a final reaction volume of 20 μL. PCR reactions were carried out in a Bio-Rad T100 Thermal Cycler using variable optimised cycling conditions. For *gatB, hcpA* and *fbpA* genes cycling was 98°C for 30 sec followed by 34 cycles of 98°C for 5 sec, 65°C for 5 sec, 72°C for 10 sec followed by 72°C for 1 min. For *coxA* and *ftsZ* genes cycling was 98°C for 30 sec followed by 34 cycles of 98°C for 5 sec, 55°C for 5 sec, 72°C for 30 sec followed by 72°C for 1 min. PCR products were separated and visualised using 2% E-Gel EX agarose gels (Invitrogen) with SYBR safe and an Invitrogen E-Gel iBase Real-Time Transilluminator.

### Sanger sequencing

PCR products were submitted to Source BioScience (Source BioScience Plc, Nottingham, UK) for PCR reaction clean-up, followed by Sanger sequencing to generate both forward and reverse reads. Where *Wolbachia* PCR primers included M13 adaptors, just the M13 primers alone (M13_adaptor_F: 5’-TGTAAAACGACGGCCAGT-3’ and M13_adaptor_R: 5’-CAGGAAACAGCTATGACC-3’) were used for sequencing, otherwise the same primers as utilised for PCR were used. Sequencing analysis was carried out in MEGAX [49]. Both chromatograms (forward and reverse traces) from each sample were manually checked, edited, and trimmed as required, followed by alignment by ClustalW and checking to produce consensus sequences. Consensus sequences were used to perform nucleotide BLAST (NCBI) database queries, and for *Wolbachia* genes searches against the *Wolbachia* MLST database (http://pubmlst.org/Wolbachia). If a sequence produced an exact match in the MLST database we assigned the appropriate allele number, otherwise we obtained a new allele number for each novel gene locus sequence for *Anopheles Wolbachia* strains through submission of the FASTA and raw trace files on the *Wolbachia* MLST website for new allele assignment and inclusion within the database. Full consensus sequences were also submitted to GenBank and assigned accession numbers. The Sanger sequencing traces from the *wsp* gene were also treated in the same way and analysed alongside the MLST gene locus scheme, as an additional marker for strain typing. Where potential mixed strains were detected (in *An. melas* and *An. gambiae* s.s-melas hybrid individuals) and any further Supergroup A or B specific testing was exhausted, it wasn’t possible to submit these sequences to the MLST database for a new allele to be assigned, however, clean 16S consensus sequences from representative individuals for each of the Supergroup A and B strains characterised were submitted to GenBank, in addition to the full MLST profile of one individual demonstrating one of the Supergroup A strain infections.

### Phylogenetic analysis

Alignments were constructed in MEGAX by ClustalW to include all relevant and available sequences highlighted through searches on the BLAST and *Wolbachia* MLST databases. Maximum Likelihood phylogenetic trees were constructed from Sanger sequences as follows. The evolutionary history was inferred by using the Maximum Likelihood method based on the Tamura-Nei model [50]. The tree with the highest log likelihood in each case is shown. The percentage of trees in which the associated taxa clustered together is shown next to the branches. Initial tree(s) for the heuristic search were obtained automatically by applying Neighbor-Join and BioNJ algorithms to a matrix of pairwise distances estimated using the Maximum Composite Likelihood (MCL) approach, and then selecting the topology with superior log likelihood value. The trees are drawn to scale, with branch lengths measured in the number of substitutions per site. Codon positions included were 1st+2nd+3rd+Noncoding. All positions containing gaps and missing data were eliminated. The phylogeny test was by Bootstrap method with 1000 replications. Evolutionary analyses were conducted in MEGAX [49].

### *Wolbachia* quantification

To estimate *Wolbachia* density across multiple mosquito species, RNA extracts were added to QubitTM RNA High Sensitivity Assays (Invitrogen) and total RNA measured using a Qubit 4 Fluorometer (Invitrogen). All RNA extracts were then diluted to produce extracts that were 2.0 nanograms (ng)/μL prior to being used in quantitative Reverse Transcription PCR (qRT-PCR) assays targeting the *Wolbachia *16S rRNA** gene [28]. A synthetic oligonucleotide standard (Integrated DNA Technologies) was designed to calculate *16S rRNA* gene copies per μL using a ten-fold serial dilution (electronic supplementary material, supplementary figure 1). *16S rRNA* gene real time qRT-PCR reactions were prepared using 5 μL of QuantiNova SYBR^®^ Green RT-PCR Kit (Qiagen), a final concentration of 1μM of each primer, 1 μL of PCR grade water and 2 μL template DNA, to a final reaction volume of 10 μL. Prepared reactions were run on a Roche LightCycler^®^ 96 System for 15 minutes at 95°C, followed by 40 cycles of 95°C for 15 seconds and 58°C for 30 seconds. Amplification was followed by a dissociation curve (95°C for 10 seconds, 65°C for 60 seconds and 97°C for 1 second) to ensure the correct target sequence was being amplified. Each mosquito RNA extract was run in triplicate alongside standard curves and no template controls (NTCs) and PCR results were analysed using the LightCycler^®^ 96 software (Roche Diagnostics).

**Figure 1.**
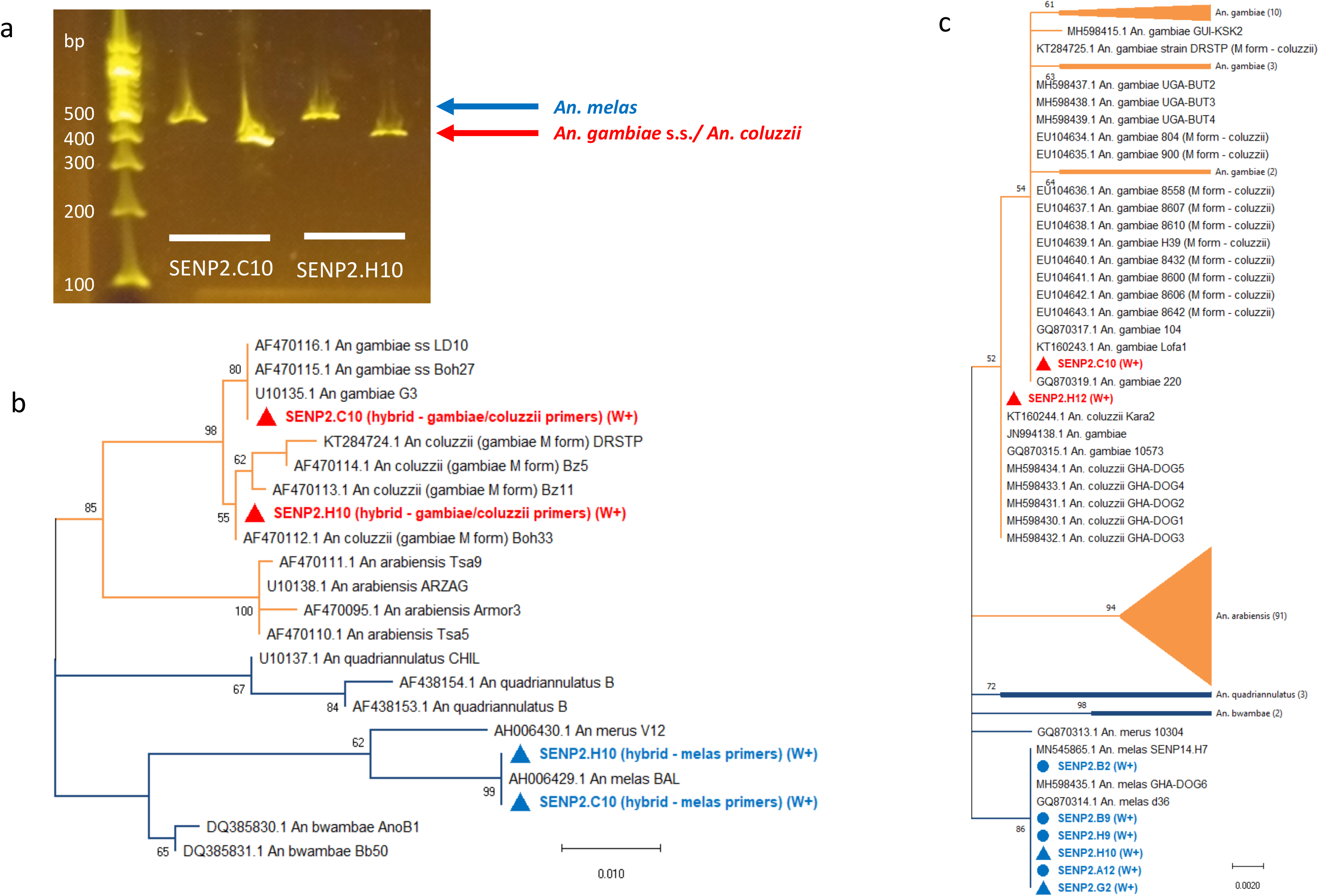
*An. gambiae* complex PCR and phylogenetic analysis of the ribosomal IGS and ITS2 gene fragments. a) Gel electrophoresis analysis of IGS *An. gambiae* / *melas* primer split down PCR products from two representative *Wolbachia* positive *An. gambiae* s.s.-*melas* hybrids. b) Maximum Likelihood molecular phylogenetic analysis of sequences from IGS gambiae / melas primer split down PCR products for representative *Wolbachia* positive (W+) hybrid samples. Sequences from IGS *An. melas* specific primer set PCR products (blue) are shown alongside IGS *An. gambiae* primer set PCR products (red). The tree with the highest log likelihood (−1966.77) is shown. The tree is drawn to scale, with branch lengths measured in the number of substitutions per site. The analysis involved 22 nucleotide sequences. There was a total of 901 positions in the final dataset. Sequences obtained from GenBank for comparison are shown with their accession numbers. c) Maximum Likelihood molecular phylogenetic analysis of *An. gambiae* complex ITS2 sequences to demonstrate ribosomal ITS2 phylogeny of *Wolbachia* positive (W+) *An. melas* (blue circles) and *An. gambiae* s.s.-*melas* hybrids (blue and red triangles). The tree with the highest log likelihood (−1360.58) is shown. The tree is drawn to scale, with branch lengths measured in the number of substitutions per site. The analysis involved 149 nucleotide sequences. There was a total of 528 positions in the final dataset. Relevant subtrees are compressed, labelled with the species and the number of sequences included within them shown in brackets. Accession numbers are shown for sequences obtained from GenBank for comparison, where they are not contained within a subtree for clear visualisation.

### *Asaia* detection

*Asaia* PCR screening was undertaken by targeting the *Asaia *16S rRNA** gene using primers Asafor: 5’-GCGCGTAGGCGGTTTACAC-3’ and Asarev: 5’-AGCGTCAGTAATGAGCCAGGTT-3’ [33, 51]. *Asaia *16S rRNA** gene real time qRT-PCR reactions were prepared using 5 μL of QuantiNova SYBR^®^ Green RT-PCR Kit (Qiagen), a final concentration of 1μM of each primer, 1 μL of PCR grade water and 2 μL template DNA, to a final reaction volume of 10 μL. Prepared reactions were run on a Roche LightCycler^®^ 96 System for 15 minutes at 95°C, followed by 40 cycles of 95°C for 15 seconds and 58°C for 30 seconds. Amplification was followed by a dissociation curve (95°C for 10 seconds, 65°C for 60 seconds and 97°C for 1 second) to ensure the correct target sequence was being amplified.

### Statistical analysis

Normalised qRT-PCR *Wolbachia *16S rRNA** gene copies per μL were compared using unpaired t-tests in GraphPad Prism 7.

## 4. Results

### Mosquito species and *Wolbachia* strain prevalence rates

In addition to confirmation of species for the morphologically indistinguishable individuals within the *An. gambiae* complex, initial screening using diagnostic species-specific PCRs highlighted the presence of some naturally occurring hybrids between members of the *An. gambiae* complex. Concomitant PCR screening demonstrated the presence of *Wolbachia* within individuals of the *An. gambiae* complex, including a number of the hybrid specimens (electronic supplementary material, table S1). The composition of these hybrids was further investigated and confirmed through repeat of the normally multiplex ribosomal IGS PCR [37] in single-plex format, separating the *An. gambiae* s.s. / *coluzzii* primer set from the *An. melas* primer set, achieving strong amplification for both target sequences (figure 1*a*) and confirmed for some representative samples through Sanger sequencing and phylogenetic analysis of both IGS PCR products from the same individuals (figure 1*b*). The further use of PCR amplification, Sanger sequencing and phylogenetic analysis of the ribosomal ITS2 (Figure 1*c*) and mitochondrial *COI, COII* (figure 2) and *ND4-ND5* (Figure 3) genes was able to confirm both the mosquito species identity for individuals of interest and the composition of hybrids, with mitochondrial gene analysis indicating the maternal species identity. Prevalence rates of natural *Wolbachia* strains were variable depending on *Anopheles* species and location (table 1). *Wolbachia* strains were detected in *An. gambiae* s.s. mosquitoes from the Faranah region with prevalence rates ranging from 0.0 - 2.8%. In the Maferinyah region, from individuals collected in Senguelen, *Wolbachia* strains were detected in *An. melas* (11.6% - 16/138) and in *An. gambiae* s.s.-*melas* hybrids (40.0% prevalence – 6/15). Interestingly, *Wolbachia* was not found in any of the 4 *An. gambiae* s.s., 18 *An. coluzzii*, 2 *An. coluzzii-gambiae* s.s. hybrids or an *An. coluzzii-melas* hybrid collected from Senguelen, suggesting *Wolbachia* strains are not currently widespread across all members of the *An. gambiae* complex in this location. Phylogenetic hybrid composition analysis combined with *Wolbachia* screening highlighted the majority of *An. gambiae* s.s.-*melas* hybrids collected from Senguelen had *An. melas* mothers (8/12 *An. melas* by mitochondrial analysis), with 4/6 *Wolbachia* positive hybrids having *An. melas* as the maternal species. These results, combined with the prevalence of maternally inherited *Wolbachia* in the *An. melas* individuals and not in the *An. gambiae* s.s. individuals from this location suggests the *Wolbachia* in this population has most likely originated from *An. melas*. *Wolbachia*-negative *An. gambiae* s.s.-*melas* hybrids were also confirmed for two specimens from Fandie (with *An. gambiae* s.s. mitochondrial results) and a *Wolbachia*-negative *An. coluzzii-gambiae* s.s. hybrid (maternally *An. coluzzii*) from Maferinyah.

**Figure 2.**
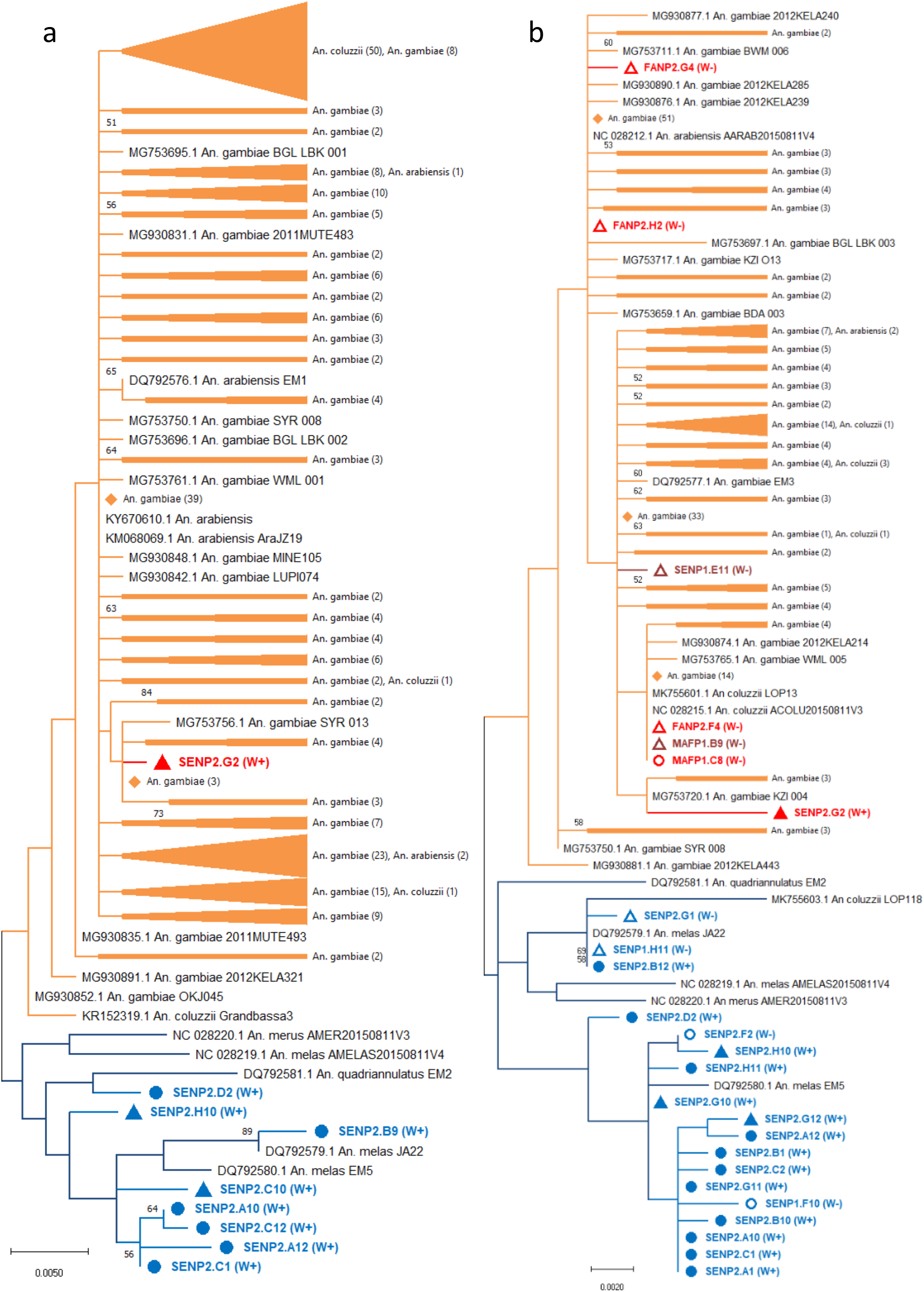
*An. gambiae* complex species phylogenetic analysis of the mitochondrial *COI* and *COII* genes. a) Maximum Likelihood molecular phylogenetic analysis of the *COI* gene for the *An. gambiae* complex. The tree with the highest log likelihood (−2546.33) is shown. The analysis involved 233 nucleotide sequences. There was a total of 658 positions in the final dataset. b) Maximum Likelihood molecular phylogenetic analysis of the COII gene for the *An. gambiae* complex. The tree with the highest log likelihood (−2013.22) is shown. The analysis involved 144 nucleotide sequences. There was a total of 728 positions in the final dataset. In both trees the branches where *An. gambiae* s.s., *An. coluzzii* and *An. arabiensis* are grouping are shown in orange, with branches where *An. melas*, *An. merus* and *An. quadriannulatus* are grouping shown in dark blue. Sequences generated in this study are shown in bold, with red for *An. gambiae* s.s., maroon for *An. coluzzii* and blue for *An. melas* sequences. Hybrid samples are shown with triangle node markers, whereas non-hybrids shown with circles. *Wolbachia* negative specimens are shown with hollow node markers and (W-) following the sample ID, whereas *Wolbachia* positives are denoted with filled node markers and (W+) following sample IDs. Accession numbers are shown for sequences obtained from GenBank for comparison, except where duplicate identical sequences were replaced with a single representative, or where subtrees are compressed, for better visualisation. One representative sequence is shown for identical duplicate sequences from the same species, labelled with the species and number of duplicate sequences in brackets, with a diamond node marker. Appropriate subtrees are compressed, labelled with the species included and the number of sequences from each respective species shown in brackets. The trees are drawn to scale, with branch lengths measured in the number of substitutions per site.

**Figure 3.**
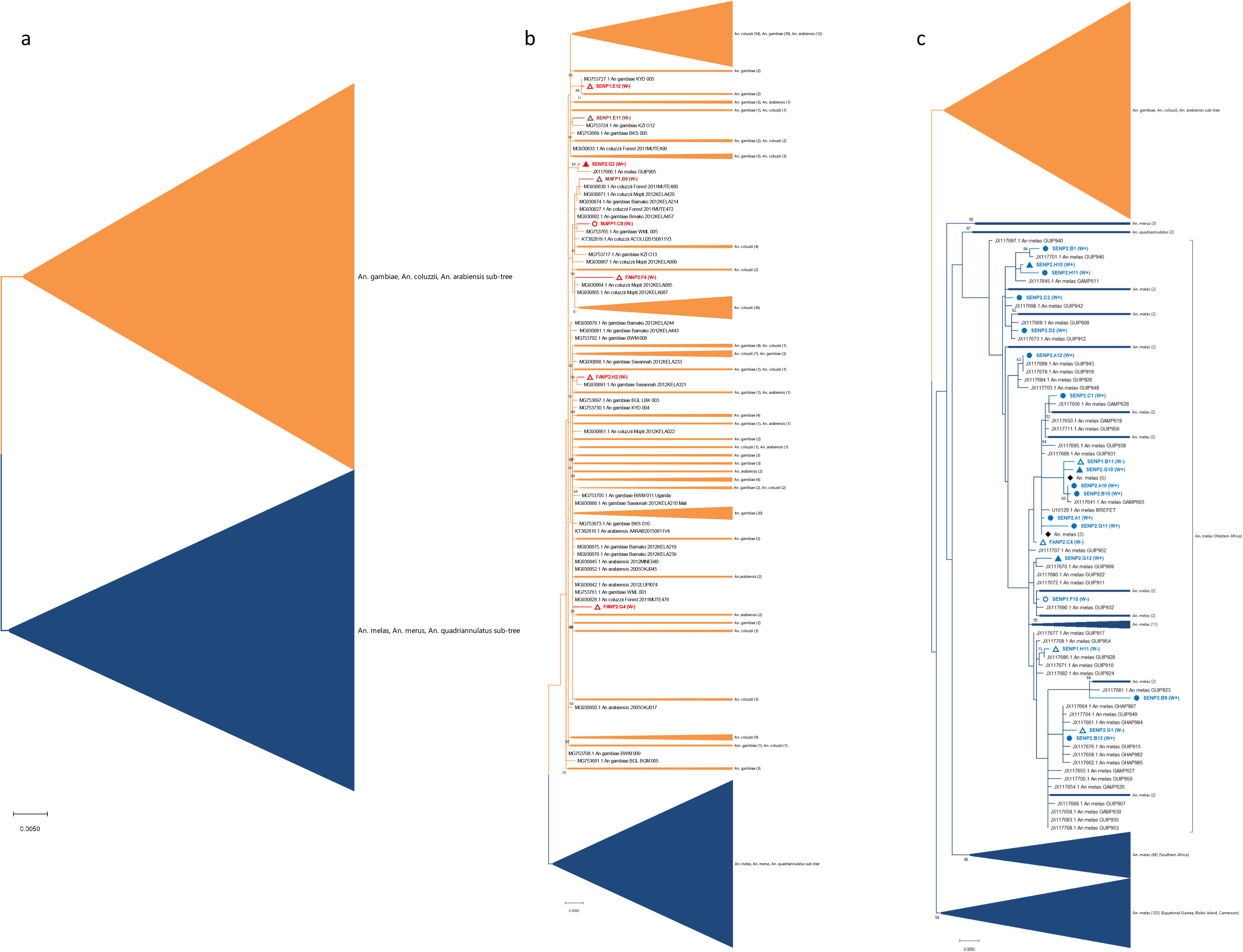
*An. gambiae* complex species phylogenetic analysis of mitochondrial *ND4-ND5* gene fragment. a) Overview of the Maximum Likelihood molecular phylogenetic analysis of the *ND4-ND5* gene fragment with a subtree compressed in orange where sequences are grouping for *An. gambiae* s.s., *An. coluzzii* and *An. arabiensis* (subtree expanded in b) and a subtree compressed in dark blue where sequences for *An. melas*, *An. merus* and *An. quadriannulatus* are grouping (subtree expanded in c). The tree with the highest log likelihood (−8134.49) is shown. The analysis involved 591 nucleotide sequences. There was a total of 1579 positions in the final dataset. Sequences generated in this study are shown in bold, with red for *An. gambiae* s.s., maroon for *An. coluzzii* and blue for *An. melas* sequences. Hybrid samples are shown with triangle node markers, whereas non-hybrids are shown with circles. *Wolbachia* negative (W-) specimens are shown with hollow node markers, whereas *Wolbachia* positives (W+) are denoted with filled node markers. Accession numbers are shown for sequences obtained from GenBank for comparison, except where duplicate identical sequences were replaced with a single representative, or where subtrees are compressed, for better visualisation. One representative sequence is shown for identical duplicate sequences from the same species, labelled with the species and number of duplicate sequences in brackets, with a diamond node marker. Appropriate subtrees are compressed, labelled with the species included and the number of sequences from each respective species shown in brackets. The tree is drawn to scale, with branch lengths measured in the number of substitutions per site. In c, square brackets denote the grouping of the Western Africa *An. melas* populations, where sequences generated in this study are also situated, with the Southern Africa and Equatorial Guinea (Bioko Island) populations clustering separately as previously found[61].

**Table 1.**
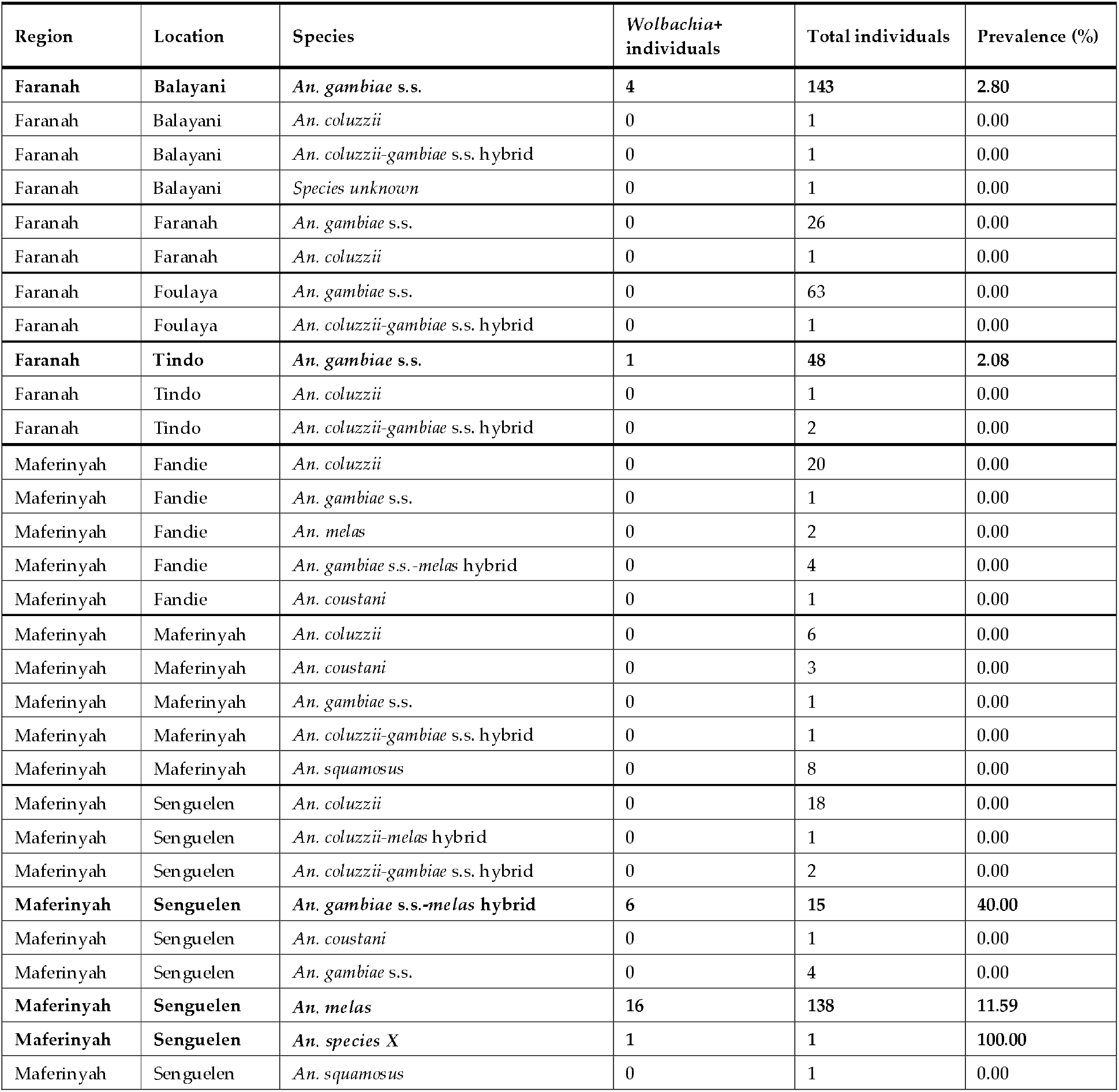
*Wolbachia* prevalence rates in *Anopheles* species collected in two regions of Guinea in 2018. Species containing *Wolbachia*-infected individuals are denoted in bold.

A *Wolbachia* strain was also found in a single female of an unclassified *Anopheles* species from Senguelen. Sanger sequencing and BLAST analysis of the ITS2 region revealed this *Anopheles* sp. ‘X’ was most similar to *Anopheles* sp. 7 BSL-2014 (GenBank accession number KJ522819.1) but at only 93.2% sequence identity, and An. theileri (GenBank accession number MH378771.1) with 90.9% sequence identity (both full query coverage). Phylogenetic analysis of the ribosomal ITS2 region and mitochondrial *COI* and *ND4-ND5* regions for An. sp. ‘X’ (Figure 4*a*, *b* and *c*) revealed that this species is from the *Myzomyia* Series, within the *Cellia* Subgenus of *Anopheles*, with agreement for this placement across all three phylogenies. The ITS2 region gave the greatest discrimination for this species, however, currently no other sequences from this species are available in order to classify it any further than to Series level and closest to, but distinct from, sequences denoted *Anopheles* sp. 7, another as yet undetermined *Anopheles* species [52]. Mosquito ribosomal and mitochondrial gene sequences were deposited in GenBank and accession numbers obtained (electronic supplementary material, table S2).

**Figure 4.**
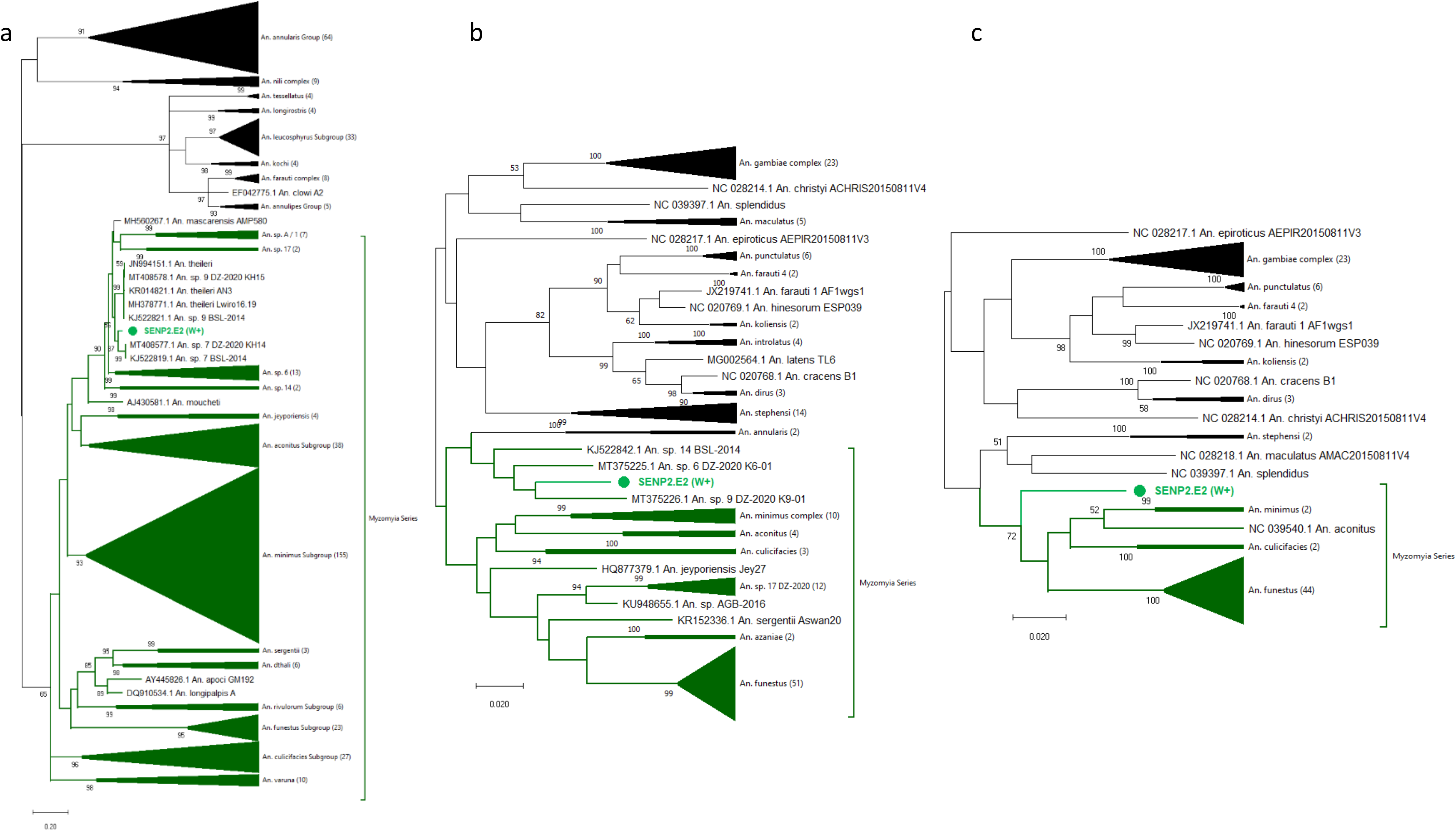
Phylogenetic analysis of the *An. sp. X* ribosomal ITS2, mitochondrial *COI* and *ND4-ND5* gene sequences within the *Cellia* Subgenus of *Anopheles*. a) Maximum Likelihood molecular phylogenetic analysis of ribosomal ITS2 sequences. All available GenBank sequences covering the sequenced fragment from the *Myzomyia, Neomyzomyia* and *Annularis* series were included, in addition to representative sequences from the *An. gambiae* complex (*Pyretophorus* Series) for broader placement and comparison with the other phylogenetic analyses. The tree with the highest log likelihood (−21743.86) is shown. The analysis involved 440 nucleotide sequences. There was a total of 1112 positions in the final dataset. b) Maximum Likelihood molecular phylogenetic analysis of mitochondrial *COI* sequences. All available GenBank sequences covering the sequenced fragment from the *Cellia* Subgenus were included. The tree with the highest log likelihood (−7291.31) is shown. The analysis involved 157 nucleotide sequences. There was a total of 658 positions in the final dataset. c) Maximum Likelihood molecular phylogenetic analysis of mitochondrial *ND4-ND5* sequences. All available GenBank sequences covering the sequenced fragment from the Cellia Subgenus were included. The tree with the highest log likelihood (−11183.21) is shown. The analysis involved 95 nucleotide sequences. There was a total of 1518 positions in the final dataset. In all trees the sequences generated in this study are shown in bold, with the *Wolbachia* positive *An. sp. X* specimen (SENP2.E2 (W+)) shown in green, with a filled circle node marker. Branches where sequences from the Myzomyia series are grouping are shown in dark green, and with an external labelled dark green bracket to denote this grouping. Relevant subtrees are compressed, labelled with the species and the number of sequences included within them shown in brackets. Accession numbers are shown for sequences obtained from GenBank for comparison, where they are not contained within a subtree for clear visualisation.

### *Wolbachia* strain typing

Although amplification of the *Wolbachia 16S rRNA* fragments of the natural strain in *An. gambiae* s.s. from the Faranah region was possible, sequences obtained were of insufficient quality for further analysis. Furthermore, no *wsp* gene amplification was possible from *An. gambiae* s.s. from the Faranah region. In contrast, *Wolbachia 16S rRNA* (figure 5) and *wsp* sequences (figure 6) were generated from both *An. melas* / *An. gambiae* s.s.-*melas* hybrids and *An. sp. X* collected from Senguelen in the Maferinyah region. Analysis of *Wolbachia 16S rRNA* sequences obtained from *An. melas* and *An. gambiae* s.s.-*melas* hybrid individuals highlighted the occurrence of superinfections within this population, with the presence of multiple *Wolbachia* strains being indicated. The *Wolbachia* 16S sequences from some *An. melas* and hybrid individuals produced consensus sequences which were most closely related to *Wolbachia* strains of Supergroup A (such as *w*Mel, *w*AlbA and *w*Au) (*w*Anga-Guinea-A), of which two different A strains (A1 and A2) could be determined in different individuals. In contrast, other *An. melas* and hybrid specimens produced *Wolbachia* 16S consensus sequences which grouped clearly with Supergroup B strains (*w*Anga-Guinea-B), also with two differing B strains able to be determined (B1 and B2) (figure 5). In addition, the sequence chromatograms from other *An. melas* and hybrid individuals consistently demonstrated mixed bases in the positions of variation between the *w*Anga-Guinea-A and *w*Anga-Guinea-B strains, with agreement both between forward and reverse sequence traces from the same individuals, as well as across multiple individuals, suggesting the presence of superinfections of both Supergroup A and B *Wolbachia* strains within these individuals.

**Figure 5.**
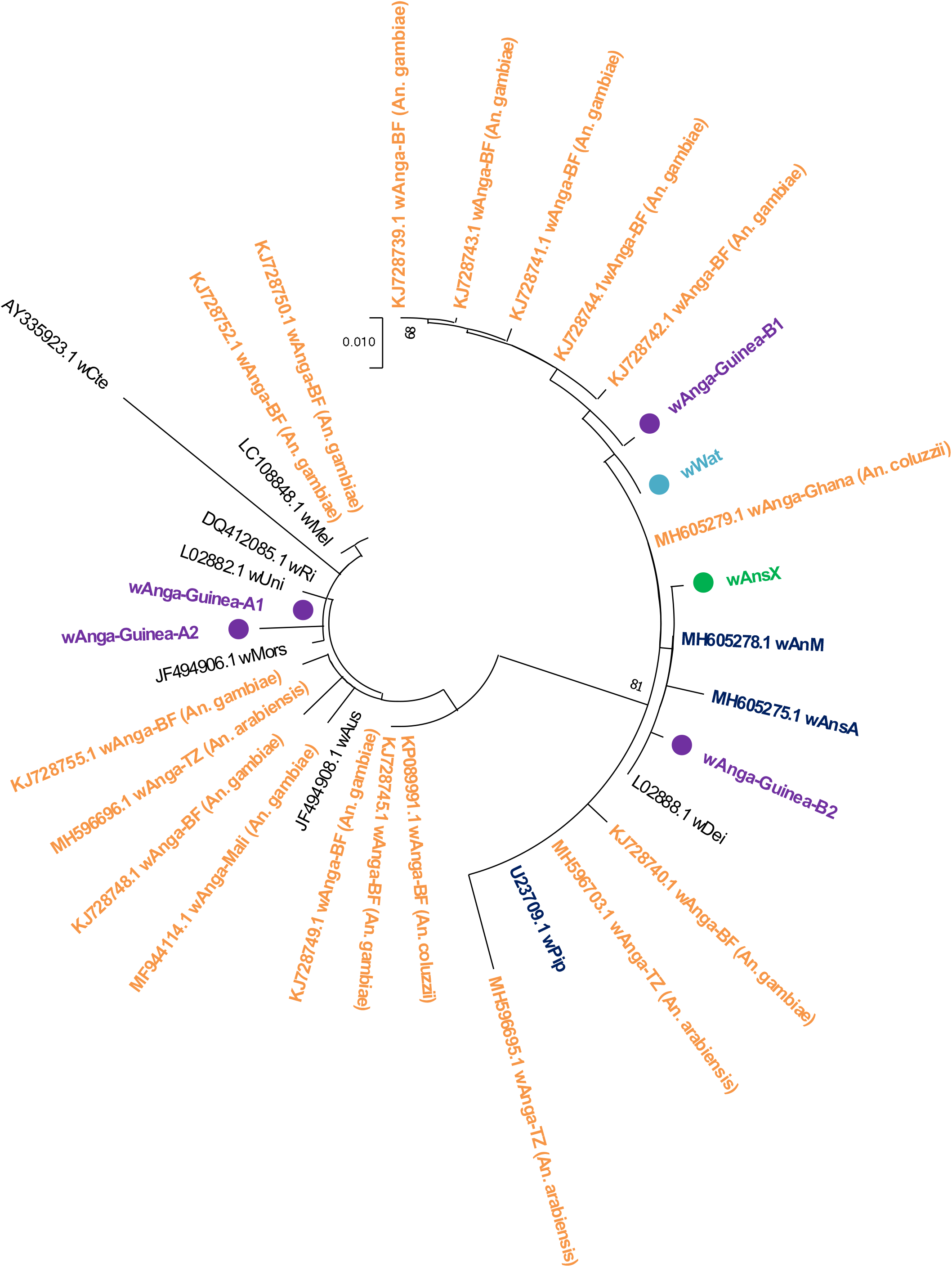
*Wolbachia* strain phylogenetic analysis using the *16S rRNA* gene. Maximum Likelihood molecular phylogenetic analysis of the *16S rRNA* gene for *Wolbachia* strains detected in *Anopheles* specimens from Guinea. The tree with the highest log likelihood (−1013.48) is shown. The tree is drawn to scale, with branch lengths measured in the number of substitutions per site. The analysis involved 34 nucleotide sequences. There was a total of 398 positions in the final dataset. Sequences generated in this study are shown in bold with filled circle node markers. The *w*Anga-Guinea sequences from both A and B Supergroups are shown in purple. The *w*AnsX strain is shown in green and the wWat strain obtained from *Cx. watti* for comparative work is shown in light blue. *w*Anga sequences obtained from the *An. gambiae* complex species previously are shown in orange with their accession numbers. *Wolbachia* strains obtained from other mosquito species in previous studies are shown in navy blue with their accession numbers. Additional *Wolbachia* sequences from non-mosquito hosts obtained from GenBank for comparison are shown with their accession numbers, in black.

**Figure 6.**
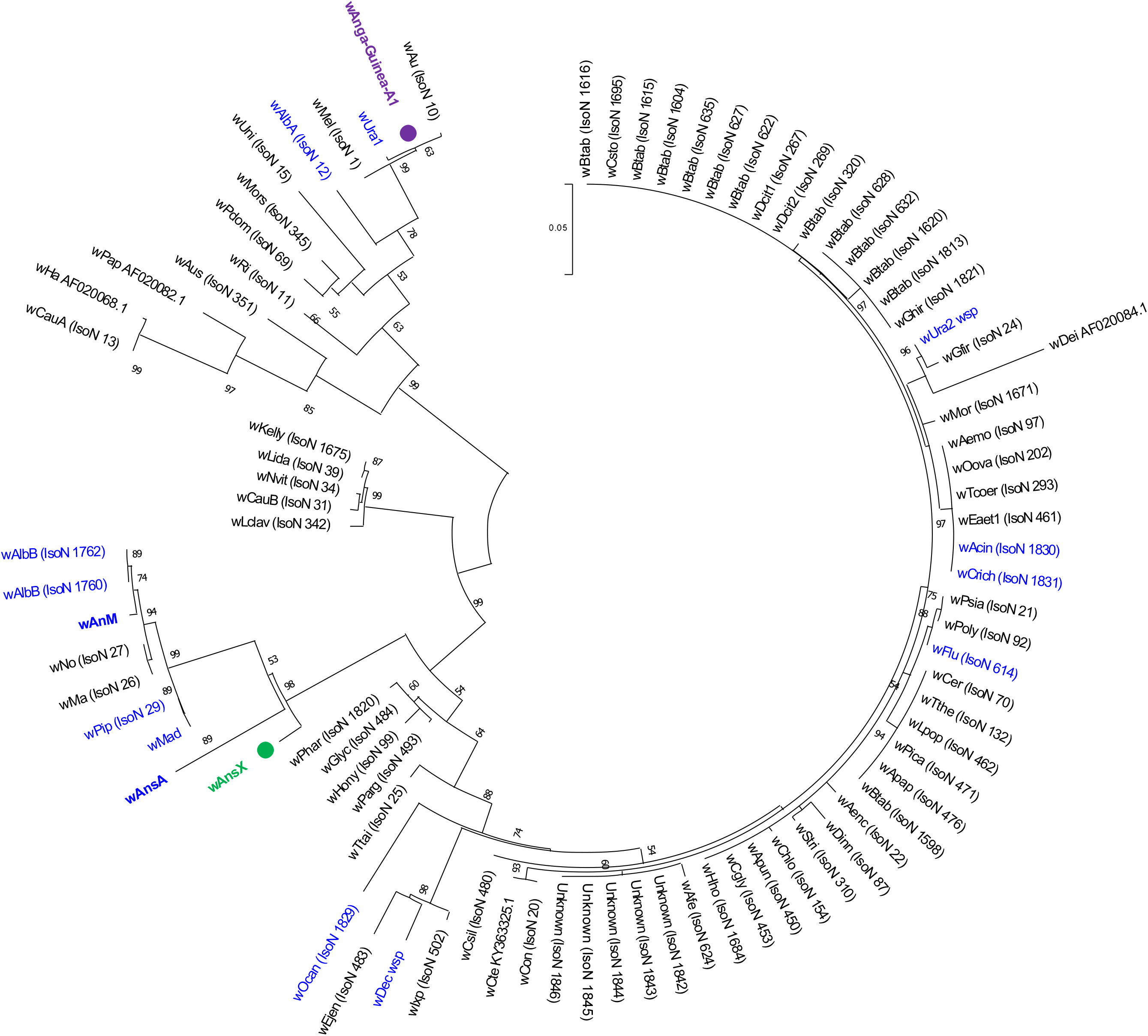
*Wolbachia* strain phylogenetic analysis using the *wsp* gene. Maximum Likelihood molecular phylogenetic analysis of the *wsp* gene for resident strains in *Anopheles* species from Guinea. The tree with the highest log likelihood (−3646.57) is shown. The tree is drawn to scale, with branch lengths measured in the number of substitutions per site. The analysis involved 86 nucleotide sequences. There was a total of 431 positions in the final dataset. Sequences generated in this study are shown in bold with filled circle node markers. *w*Anga-Guinea-A1 *wsp* from an *An. melas* specimen is shown in purple, and *w*AnsX is shown in green. Reference numbers of additional sequences obtained from the MLST database (IsoN; Isolate number) or GenBank (accession number) are shown. Strains isolated from mosquitoes are shown in blue, with those strains from other *Anopheles* species highlighted in bold.

To try to further understand the complex picture of *Wolbachia* strains in the *An. melas* and *An. gambiae* s.s.-*melas* hybrids from Senguelen, certain *An. melas* and hybrid individuals were selected for repeat *Wolbachia 16S* sequencing, including representative samples where the original analysis suggested the presence of a *w*Anga-Guinea A and B strain superinfection and where there was apparent dominance of one or other strain. Comparison of the repeat 16S sequences to the original analysis for each individual suggested that the presence of superinfections was genuinely evident but that it wasn’t possible to confidently separate *An. melas* and hybrid individuals into *w*Anga-Guinea-A only, *w*Anga-Guinea-B only, or superinfected groups, with the dominance of a particular strain over another, on the basis of 16S sequencing analysis alone. This was due to variation in the dominance of *w*Anga-Guinea-A or *w*Anga-Guinea-B sequences in chromatograms, suggesting the possibility that any difference could be due to normal technical variations in the processes of amplification and sequencing (e.g. the apparent dominance of either an A or B strain in chromatograms from a particular PCR product could be due to the chance of variation in amplification efficiency and the resultant sequencing signal strength), rather than detection of a repeatable biological difference, with consistent dominance of a particular strain variant over another within each individual. This repeat analysis on a small sub-group suggested that *Wolbachia* superinfections were likely in all the *An. melas* or hybrid individuals where this subsequent *16S* analysis was possible.

This, combined with the overall results from the *Wolbachia 16S* analysis from all infected individuals suggested superinfections were widespread in *Wolbachia* positive individuals but there did not currently appear to be a clear dominant strain, or strain variant, which could be identified with greater relative occurrence or apparent density (through consistent stronger sequencing signal strength) to the other strain(s) present in individuals from this population. This complexity was also mirrored when looking between *An. melas* and the hybrid specimens, with no clear distinction in the *Wolbachia* strain variants apparent in each group. Unfortunately, further comparative analysis of differing strains in *An. melas* and hybrid individuals utilising the *Wolbachia wsp* gene locus was not possible, as *wsp* sequence could only be successfully obtained from one *An. melas* individual, where *16S* analysis had indicated the presence of *w*Anga-Guinea-A1 only. This *wsp* sequence matched allele 23 within the *Wolbachia* MLST database (table 2), demonstrating that it is identical to the *wsp* sequences obtained from 20 other Supergroup A *Wolbachia* isolates contained within the database.

**Table 2.** Novel resident *Wolbachia* strain *WSP* typing and multilocus sequence typing (MLST) gene allelic profiles. Newly assigned novel alleles for *w*AnsX are shown in bold red font. \**w*Anga-Guinea-A1 *hcp*A could not be assigned a novel allele number due to a possible double infection which was unresolvable, therefore the allele number of the closest match (CM) is shown with the number of single nucleotide differences to the closest match in brackets.

| Mosquito species | Wolbachia strain | WSP typing allele numbers |  |  |  |  | MLST gene allele numbers |  |  |  |  |
| --- | --- | --- | --- | --- | --- | --- | --- | --- | --- | --- | --- |
|  |  | *wsp* | HVR1 | HVR2 | HVR3 | HVR4 | *gatB* | *coxA* | *hcpA* | *ftsZ* | *fbpA* |
| *An. melas* | *w*Anga-Guinea-A1 | 23 | 1 | 12 | 21 | 19 | 1 | 1 | CM1 (2)* | 3 | 1 |
| *An. sp. X* | *w*AnsX | 737 | 264 | 297 | 3 | 323 | 285 | 282 | 310 | 246 | 454 |

Phylogenetic analysis of both *Wolbachia 16S* and *wsp* gene fragments from *An. sp*. X indicated that the *w*AnsX strain is most closely related to *Wolbachia* strains of Supergroup B (such as *w*Pip, *w*AlbB, *w*AnsA, *w*AnM, *w*Ma and *w*No). Typing of the *w*AnsX *wsp* nucleotide sequence highlighted that there were no exact matches to *wsp* alleles currently in the *Wolbachia* MLST database (https://pubmlst.org/Wolbachia/), and only one of the four hypervariable regions (HVRs) matched a known sequence (HVR3: allele 3). All *Wolbachia* gene sequences of sufficient quality to generate a consensus were deposited into GenBank and accession numbers obtained (electronic supplementary material, supplementary tables S3 and S4).

*Wolbachia* MLST was undertaken to attempt to provide more accurate strain discrimination and phylogenies. This was successfully done for the novel *Anopheles Wolbachia* strains *w*Anga-Guinea-A1 and *w*AnsX but no amplification was seen for any of the five MLST genes from *Wolbachia*-infected *An. gambiae* s.s. from Faranah, and successful sequencing of complete MLST profiles of sufficient quality for onward analysis was not possible from any further *Wolbachia* positive *An. melas* or *An. gambiae* s.s.-*melas* hybrid individuals from Senguelen. MLST gene fragment amplification was variable for *w*Anga-Guinea strains found in *An. melas* and *An. gambiae* s.s.-*melas* hybrids. Even for *w*Anga-Guinea-A1, the use of *hcpA* ‘A strain specific’ primers (hcpA_F1: GAAATARCAGTTGCTGCAAA, hcpA_AspecR1: TTCTARYTCTTCAACCAATGC) was required to generate sequence of sufficient quality for analysis of the *hcpA* gene and to therefore complete the *Wolbachia* MLST profile for this *An. melas* sample. Despite this use of Supergroup specific primers, there was still some indication of a possible mixed strain from the chromatograms generated from sequencing the A strain-specific hcpA PCR product, through mixed bases at two positions on both the forward and reverse reads, although a consensus could still be obtained through agreement of the strongest base at each position across reads. The resultant MLST allelic profile for *w*Anga-Guinea-A1 (table 2) was closest to the profile for Strain Type 13, with the variation occurring in the two positions of mixed bases in the *hcpA* locus, being closest to *hcpA* allele 1, except for a change from G to A at position 313 and from A to G at position 319 on this locus (table 2). This may indicate the presence of both *hcpA* allele 1 and an *hcpA* variant. Even if *w*Anga-Guinea-A1 were identical to Strain Type 13, of the 19 records available on the MLST database (all Supergroup A), no other isolates with this strain type, where host information had been provided, were found in mosquito species. Concatenation of the MLST loci and phylogenetic analysis also confirms *w*Anga-Guinea-A1 is closest to strains belonging to Supergroup A, including *w*Mel and *w*AlbA (as also suggested by *16S* and *wsp* gene phylogenies). For *w*AnsX, new alleles for all five MLST gene loci (sequences differed from those currently present in the MLST database), and the therefore novel allelic profile, confirms the diversity of this novel *Wolbachia* strain (table 2). The phylogeny of *w*AnsX based on concatenated sequences of all five MLST gene loci confirms this strain clusters within Supergroup B and further demonstrates that it is distinct from other currently available strain profiles (figure 7). Consistent with previous studies looking at novel *Wolbachia* strains in *Anopheles* species using MLST [24], these results highlight the lack of concordance between *Wolbachia* strain phylogeny and their insect hosts across diverse geographical regions.

**Figure 7.**
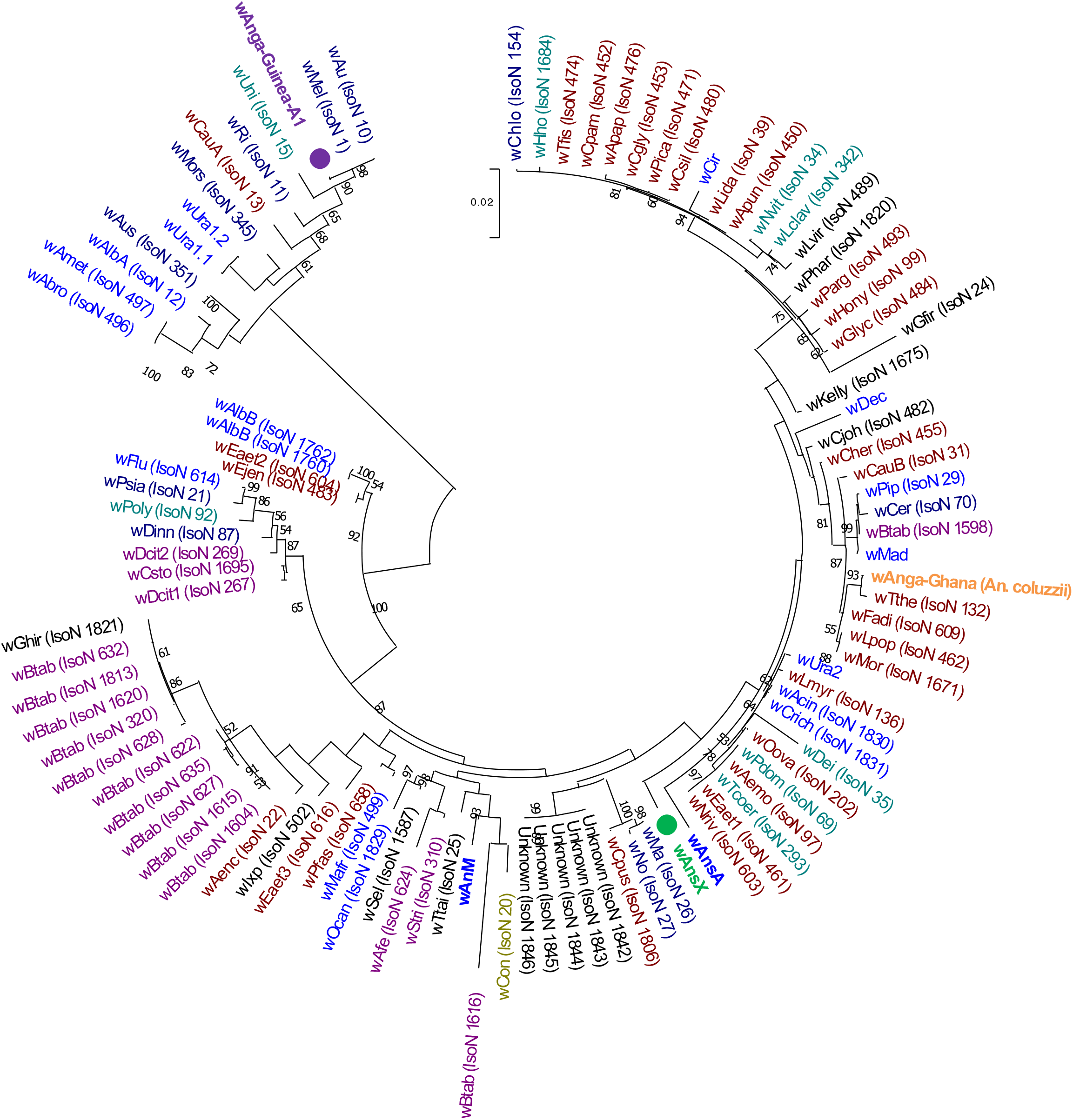
*Wolbachia* multilocus sequence typing (MLST) phylogenetic analysis of *w*Anga-Guinea-A1 and *w*AnsX. Maximum Likelihood molecular phylogenetic analysis from concatenation of all five MLST gene loci for resident *Wolbachia* strains *w*Anga-Guinea-A1 (in purple) and *w*AnsX (in green). Concatenated sequences obtained in this study are highlighted in bold with a filled circle node marker. The tree with the highest log likelihood (−11404.41) is shown and drawn to scale, with branch lengths measured in the number of substitutions per site. The analysis involved 102 nucleotide sequences. There were a total of 2063 positions in the final dataset. The concatenated MLST sequence data from *w*Anga-Ghana obtained from *An. coluzzii* in a previous study[26] is shown in orange. Concatenated sequence data from *Wolbachia* strains downloaded from the MLST database for comparison are shown with isolate numbers in brackets (IsoN). *Wolbachia* strains isolated from mosquito species are shown in blue, with those strains from other *Anopheles* species highlighted in bold. Strains isolated from other Dipteran species are shown in navy blue, from Coleoptera in olive green, from Hemiptera in purple, from Hymenoptera in teal blue, from Lepidoptera in maroon and from other, or unknown orders in black.

### *Wolbachia* strain densities and relative abundance

The relative densities of *Wolbachia* strains were estimated using qRT-PCR targeting the *16S rRNA* gene after first standardising total RNA (ng per reaction). This allowed direct comparisons between phylogenetically diverse *Anopheles* species and accounts for variation in mosquito body size and RNA extraction efficiency between samples. This also allows a comparison to another novel natural *Wolbachia* strain present in *Cx. watti* (termed wWat strain) collected in Maferinyah, contemporaneously with the *Anopheles* specimens. *16S rRNA* qRT-PCR analysis revealed a mean of 1.50E+04 (+/− 4.37E+03) *16S rRNA* copies/μL for the *w*AnsX strain in the single individual (figure 8, electronic supplementary material, supplementary table S5). Lower mean densities were found for the *w*Anga-Guinea strains in *An. melas* individuals (n=14) and *An. gambiae* s.s.-*melas* hybrids (n=4) with 8.20E+02 (+/−2.90E+02) and 1.41E+02 (+/− 3.95E+01) *16S rRNA* copies/μL respectively. The densities were compared to the wWat strain in *Cx. watti* females also collected in the Maferinyah region with a mean density of 2.37E+04 (+/− 5.99E+03). The density of the wWat strain was significantly higher than the *w*Anga-Guinea strains found in *An. melas* and hybrids (Unpaired T-test, p=0.002). Individual *An. gambiae* s.s. extracts from the Faranah region that were identified as *Wolbachia*-infected by amplification of the *16S rRNA* gene [44] did not result in any *16S rRNA* qRT-PCR amplification, suggesting a very low titre *Wolbachia* strain present in these individuals.

**Figure 8.**
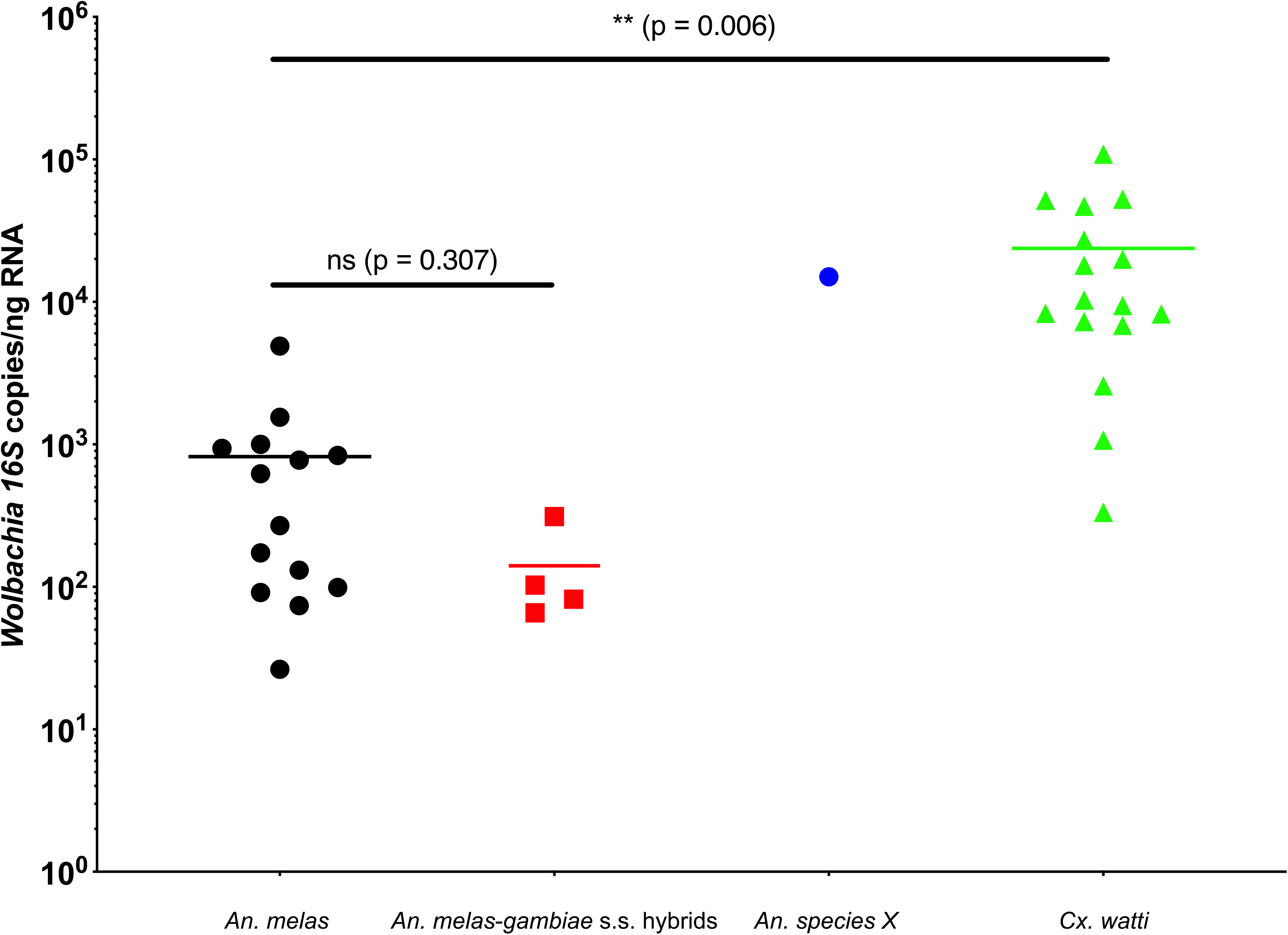
*Wolbachia* strain densities in wild-caught female mosquitoes from the Maferinyah sub-prefecture of Guinea. Total RNA extracted from individual mosquitoes was standardised to 2.0 ng/μL prior to being used in qRT-PCR assays targeting the *Wolbachia *16S rRNA** gene. A synthetic oligonucleotide standard was designed to calculate *16S rRNA* gene copies per μL of RNA using a serial dilution series and all samples were run in triplicate in addition to no template controls.

### *Wolbachia* and *Asaia* co-infections

Individual mosquitoes shown to be infected with the *w*AnsX or *w*Anga-Guinea strains were screened for the presence of *Asaia* bacteria using qRT-PCR. Co-infections were detected in all *An. melas* (n=14, mean *Asaia 16S rRNA* Ct value = 30.60 +/− 2.02), all *An. gambiae* s.s.-*melas* hybrids (n=4, mean *Asaia 16S rRNA* Ct value =26.32 +/−3.54) and in the single *An. species* X (*Asaia 16S rRNA* Ct value = 34.92) (electronic supplementary material, supplementary table S5).

## 5. Discussion

Endosymbiotic *Wolbachia* bacteria are particularly widespread through insect populations but were historically considered absent from the *Anopheles* genera [19]. The discovery of additional novel natural strains of *Wolbachia* in *Anopheles* species suggests that the prevalence and diversity has been significantly under-reported to date. Since 2014, there have been several reports of detection of *Wolbachia* strains in major malaria vectors, such as sibling species in the *An. gambiae* complex [25, 26, 28-30] and *An. moucheti* [26]. This study provides evidence for *Wolbachia* strains in *An. melas*, a species within the *An. gambiae* complex, which can be an important local vector of malaria in West-African coastal areas where it breeds in brackish water, mangrove forests and salt marshes [53, 54]. It’s importance as a local malaria vector was shown in Equatorial Guinea where the average number of malaria infective *An. melas* bites/person/year was recorded at up to 130 [55]. The finding of natural *An. gambiae* s.s.-*melas* hybrids in this study appears highly unusual, with published accounts of hybridisation between members of the *An. gambiae* complex seeming to agree that detection of hybrids in wild populations is relatively rare [56], and when it does occur, seems most often to be a combination of hybrids between *An. gambiae* s.s., *An. coluzzii* or *An. arabiensis*. Historical reports of *An. gambiae* s.s.-*melas* hybrids were also in West Africa but with laboratory colonies, giving variable results for ongoing success of hybrid colonies [57–59]. Interestingly colonised *An. melas* and F1 hybrid larvae were able to be reared in distilled water in the laboratory, rather than requiring a higher salinity content as might be expected from the natural ecology of *An. melas* [59]. As *An. melas* is more geographically restrained and has a more defined ecological niche than other members of the *An. gambiae* complex, natural hybrids composed of these constituent species are arguably less likely to occur, with fewer areas of sympatry. Natural hybrids may also be underestimated [60] due to sampling bias with a greater proportion of studies focussing on the more widely distributed major anthrophilic malaria vectors, *An. gambiae* and *An. arabiensis* [61].

Hybrid detection is also dependent on the methodology used for species identification and the format of species-specific diagnostic assays [37, 56]. Our testing highlighted that amplification and clarity of hybrid detection was improved with use of the ribosomal IGS PCR primers[37] for each species in single-plex format, rather than the standard higher-throughput multiplex format, where primers for multiple members of the *An. gambiae* complex are included at the same time, with different product sizes for species discrimination. This is unsurprising due to the designed aims of the multiplex assay, and potential variations in reaction efficiency between species, particularly when hybridised, which were highlighted in the original publication [37]. However, this could potentially result in reduced detection of natural hybrids, compared to the apparent detection of individual species, when used for widespread screening and species identification. Sanger sequencing of the single-plex species-specific IGS PCR products for representative hybrid samples enabled confirmation of the hybridisation and the avoidance of doubt from any possibility of specificity problems [37, 56], before further confirmation was obtained through subsequent sequencing and phylogenetic analysis of other gene fragments.

Genetic divergence also likely affects interspecific hybrids and the original delineation of the member species within the *An. gambiae* complex was concluded on the basis of hybrid male sterility from early crossing experiments [61]. However, the full extent and impacts of interspecific hybridisation between members of the complex is still under investigation and debate [60]. *An. gambiae* s.s. and *An. melas* have a greater degree of genetic divergence from one another when compared to other members of the complex (such as *An. gambiae* s.s., *An. coluzzii* and *An. arabiensis*) and *An. melas* groups separately and more closely to *An. merus* and *An. quadriannulatus* sequences. Even within *An. melas*, species-specific microsatellite markers and mitochondrial genetic analysis of geographically distinct populations suggested there was species level divergence between different populations, resulting in three distinct major clusters; Bioko Island, Western mainland and Southern mainland African populations (with mainland population division occurring in Cameroon) [61]. In the context of the results of this study, our *An. melas* would be included in the Western mainland cluster (this is supported by our phylogenetic analysis). Following the discovery of *Wolbachia* in this population, it would be interesting to investigate whether *Wolbachia* strains were also present in other *An. melas* geographic clusters, and whether the CI phenotype was evident in some or all of these strains. If stable *Wolbachia* infections were present in some populations but not others, it also raises the question of the length of time *Wolbachia* may have been present in this species and whether *Wolbachia* infections may be having an influence on the host population genetics and affecting genetic divergence and speciation over time.

The discovery of the *w*AnsX strain led to retrospective confirmation of the host mosquito species using Sanger sequencing. In this study and previous studies, thorough and accurate molecular identification is important given the difficulties of morphological identification, the potential for currently unrecognised cryptic species [52, 62] and potential for inaccuracies for certain species where only diagnostic species PCR-based methods are used for molecular identification [63]. Phylogenetic analysis and confident species discrimination is dependent on the sequences available for comparison at the time. Sequencing and phylogenetic analysis of all three regions for this specimen indicated placement within the *Cellia* Subgenus and *Myzomyia* Series of *Anopheles*, with the greatest number of closely related comparative sequences available for comparison in the ITS2 region. Our analysis revealed that this species is closest to *Anopheles* sp. 7, followed by An. theileri from sequences currently available. *Anopheles* sp. 7 BSL-2014 was collected in the Western Kenyan Highlands, with 1 of 23 specimens Plasmodium falciparum ELISA sporozoite and PCR positive[52]. *An. theileri* was collected in the Democratic Republic of Congo [64] and was found to be infected with *Plasmodium sporozoites* in eastern Zambia [65].

The results of this study also highlight the requirement to provide as much genetic information and confirmation as possible for a newly discovered strain of *Wolbachia* (particularly low-density infections). The first discovery of *Wolbachia* strains in wild *An. gambiae* populations in Burkina Faso resulted from sequencing of the *16S rRNA* gene rather than screening using *Wolbachia*-specific genes [25]. A more recent comprehensive analysis through screening of *An. gambiae* genomes (Ag1000G project) concluded that determining whether a *Wolbachia* strain is present in a given host based on the sequencing of one gene fragment (often *16S rRNA*) is problematic and caution should be taken [31]. In this study, we were only able to amplify a *Wolbachia 16S rRNA* gene fragment from *An. gambiae* s.s., which is consistent with numerous recent studies in which low density strains have been detected [27, 30]. As a result, caution must be taken in drawing conclusions on the stability of infection and biological significance. Other explanations for the amplification of *16S rRNA* gene fragments include *Wolbachia* DNA insertions into an insect chromosome or contamination from non-mosquito material such as ectoparasites or plants [31]. In contrast to previous studies, we extracted RNA increasing the chances that detection of the *16S rRNA* gene is from actively expressed *Wolbachia* and indicating amplification is more likely of bacterial gene origin (rather than through integration into the host genome). However, these results are consistent with previous studies in which every *Wolbachia 16S rRNA* amplicon and sequence attributed to *An. gambiae* s.s. is unique and appears at very low density [31].

The densities of the *w*Anga-Guinea and *w*AnsX strains detected in Senguelen (measured using qRT-PCR) are significantly higher than *Wolbachia* detected in *An. gambiae* s.s. from Faranah (which were not detectable using this qRT-PCR assay targeting the *16S rRNA* gene). The *w*Anga-Guinea strains appear to have both an intermediate prevalence rate and density and further studies are required to elucidate the relative density contribution and possible differential localisation of these *Wolbachia* strains within the mosquito host, whether these strains may be influencing host population genetics (including the occurrence of natural hybrids and the intraspecific diversity within *An. melas*) and investigate these strains across more diverse geographical areas. Caution and further investigation is also required for the *w*AnsX strain as this was detected from the only collected individual of this unclassified *Anopheles* species. The detection of *Wolbachia*-*Asaia* co-infections in all individuals was in contrast to our previous study [26] but *Asaia* can be environmentally acquired at different mosquito life stages and the prevalence and density was significantly variable across different *Anopheles* species and locations [26]. These contrasting results suggest a complex association between these two bacterial species in wild *Anopheles* mosquito populations and given that *Asaia* is environmentally acquired, this association will be highly location dependent.

*Wolbachia* strains in *An. species* A (*w*AnsA) and *An. moucheti* (*w*AnM) [26], and now *An. melas* (*w*Anga-Guinea-A1) and *An. sp. X* (*w*AnsX), have complete MLST and *wsp* profiles and are at significantly higher densities when compared to strains detected in *An. gambiae* s.s. from the same countries. As *Wolbachia* density is strongly correlated with arbovirus inhibition in Aedes mosquitoes [5, 7, 11, 12], higher density strains in *Anopheles* species would be predicted to have a greater impact on malaria transmission in field populations. In this study, we screened for *P. falciparum* infection and found very low prevalence rates (<1%; data not shown) preventing any statistical analysis on *Wolbachia-Plasmodium* interactions. This study and previous studies measuring a direct impact on *Plasmodium* infection in wild populations are dependent on parasite infection rates which can be low even in malaria-endemic areas [26] and particularly for the infective sporozoite stage [66]. Low pathogen prevalence rates are also limiting factors in assessing the effect of natural strains of *Wolbachia* on arboviruses in wild mosquito populations [67]. In addition to looking at effects on *Plasmodium* prevalence in field populations, further work should look to undertake vector competence experiments with colonised populations and to determine if these *Wolbachia* strains are present in tissues such as the midgut and salivary glands which are critical to sporogony. Further studies are also needed to determine if the *w*Anga-Guinea strains are maternally transmitted given our results would suggest they are likely to be from the *An. melas*, rather than from *An. gambiae* s.s. Furthermore, an assessment of how these *Wolbachia* strains are being maintained in field populations is needed, and to determine if the CI reproductive phenotype can be induced by these strains (and if it affects viability of subsequent generations). As the chances of success of *Wolbachia* transinfection experiments can be improved by the adaptation of the *Wolbachia* strain to the target host genetic background[68], this may imply favourable potential for *w*Anga-Guinea transinfection experiments and successful establishment of a stable *Wolbachia* infection within *An. gambiae* s.s. colonies, in addition to An. melas. If achievable, this would be a big step forward in determining whether these strains (which appear relatively higher density than *Wolbachia* previously detected in the *An. gambiae* complex) could reduce malaria transmission through *Wolbachia*-based biocontrol strategies.

## 6. Conclusion

Although the debate continues over the biological significance (or even presence of natural strains in the *An. gambiae* complex), this study provides strong evidence of additional novel strains with relatively higher density infections, in addition to *Wolbachia* positive natural hybrids in the *An. gambiae* complex, and may reflect the under-reporting of natural strains in the *Anopheles* genus. The presence of *Wolbachia* superinfections increases the complexity of phylogenetic characterisation of individual strains and the determination of the relative contribution of each strain to the overall density. There are previous studies showing natural *Wolbachia* superinfections in wild mosquito populations such as *Ae. albopictus* [69] and superinfections have been generated in mosquitoes used for biocontrol strategies [7, 70] indicating superinfections can form stable associations with mosquito hosts. Candidate *Wolbachia* strains for mosquito biocontrol strategies require synergistic phenotypic effects to impact the transmission of mosquito-borne pathogens and further studies are needed to determine if these strains would induce CI and what effects they may have on host fitness. Whether these *Wolbachia* superinfections can inhibit Plasmodium parasites [28, 29] or influence the ability to transinfect other *Wolbachia* strains for population suppression and replacement strategies [71] remains to be determined but further investigation is warranted.

## Acknowledgments

The following people are thanked for their efforts in the original collection of mosquitoes: Moussa Sylla, Gnepou Camara, Louisa Messenger, Patrick Heard, Yaya Barry, Denka Camara, and Ismael Yansane. This publication made use of the PubMLST website (https://pubmlst.org/Wolbachia/) sited at the University of Oxford (<u>Jolley & Maiden 2010, BMC Bioinformatics, 11:595</u>). The development and maintenance of this site has been funded by the Wellcome Trust.

## Ethical Statement

Mosquito collection protocols were reviewed and approved by the Comite National d’Ethique pour la Recherche en Sante (030/CNERS/17) and the institutional review boards (IRB) of the London School of Hygiene and Tropical Medicine (#14798 and #15127) and the Centers for Disease Control and Prevention, USA (2018-086); all study procedures were performed in accordance with relevant guidelines and regulations. Fieldworkers participating in human landing catches were provided with malaria prophylaxis for the duration of the study.

## Funding Statement

CLJ and TW were supported by Wellcome Trust /Royal Society grants awarded to TW (101285/Z/13/Z): http://www.wellcome.ac.uk; https://royalsociety.org. SRI was supported by the President’s Malaria Initiative (PMI)/CDC.

## Data Accessibility

The datasets supporting this article have been uploaded as part of the Supplementary Material and all the sequencing data generated is available in GenBank with accession numbers as shown in the relevant supplementary tables.

## Competing interests

*We have no competing interests*.

## Authors’ Contributions

C.L.J. designed the study, carried out the analyses and drafted the manuscript. C.C-U. performed field work, carried out preliminary laboratory analysis and reviewed the manuscript. A.H.B. supervised field work and reviewed the manuscript. C.S. performed field work and reviewed the manuscript. E.K.L. supervised fieldwork and reviewed the manuscript. M.K. performed field work and reviewed the manuscript. S.R.L. supervised fieldwork and reviewed the manuscript. T.W. designed the study, carried out analyses, drafted the manuscript and provided overall supervision.

## Notes

### Competing Interest Statement

The authors have declared no competing interest.

### Summary of Updates

Wolbachia superinfections present in hybrids between An. gambiae ss and An. melas.

