## Supplementary Table 2 for "Evidence for natural hybridisation and novel *Wolbachia* strain superinfections in the *Anopheles gambiae* complex from Guinea"

| **Sample ID** | **Sample species** | **Gene fragment** | **Sequence identity** | **Accession number** |
| --- | --- | --- | --- | --- |
| SENP2.C10 | *An. melas-gambiae* s.s. hybrid | Scott IGS gambiae | *An. gambiae* s.s. | MW179559 |
| SENP2.H10 | *An. melas-gambiae* s.s. hybrid | Scott IGS gambiae | *An. gambiae* s.s. | MW179560 |
| SENP2.C10 | *An. melas-gambiae* s.s. hybrid | Scott IGS melas | *An. melas* | MW179561 |
| SENP2.H10 | *An. melas-gambiae* s.s. hybrid | Scott IGS melas | *An. melas* | MW179562 |
| SENP2.B2 | *An. melas* | ITS2 | *An. melas* | MW172421 |
| SENP2.G2 | *An. gambiae* s.s.*-melas* hybrid | ITS2 | *An. melas* | MW172422 |
| SENP2.B9 | *An. melas* | ITS2 | *An. melas* | MW172423 |
| SENP2.H9 | *An. melas* | ITS2 | *An. melas* | MW172424 |
| SENP2.C10 | *An. melas-gambiae* s.s. hybrid | ITS2 | *An. gambiae* s.s. | MW172425 |
| SENP2.H10 | *An. melas-gambiae* s.s. hybrid | ITS2 | *An. melas* | MW172426 |
| SENP2.A12 | *An. melas* | ITS2 | *An. melas* | MW172427 |
| SENP2.H12 | *An. gambiae* s.s.*-melas* hybrid | ITS2 | *An. gambiae* s.s. | MW172428 |
| SENP2.E2 | *An. species X* | ITS2 | *An. species X* | MW172429 |
| SENP2.C1 | *An. melas* | Folmer COI | *An. melas* | MW168822 |
| SENP2.D2 | *An. melas* | Folmer COI | *An. melas* | MW168823 |
| SENP2.G2 | *An. gambiae* s.s.*-melas* hybrid | Folmer COI | *An. gambiae* s.s. | MW168824 |
| SENP2.B9 | *An. melas* | Folmer COI | *An. melas* | MW168825 |
| SENP2.A10 | *An. melas* | Folmer COI | *An. melas* | MW168826 |
| SENP2.C10 | *An. melas-gambiae* s.s. hybrid | Folmer COI | *An. melas* | MW168827 |
| SENP2.H10 | *An. melas-gambiae* s.s. hybrid | Folmer COI | *An. melas* | MW168828 |
| SENP2.A12 | *An. melas* | Folmer COI | *An. melas* | MW168829 |
| SENP2.C12 | *An. melas* | Folmer COI | *An. melas* | MW168830 |
| SENP2.E2 | *An. species X* | Folmer COI | *An. species X* | MW168831 |
| SENY67 | *Cx. watti* | Kumar *CO1* | *Cx. watti* | MW168832 |
| SENP2.A1 | *An. melas* | COII | *An. melas* | MW179563 |
| SENP2.B1 | *An. melas* | COII | *An. melas* | MW179564 |
| SENP2.C1 | *An. melas* | COII | *An. melas* | MW179565 |
| SENP2.C2 | *An. melas* | COII | *An. melas* | MW179566 |
| SENP2.D2 | *An. melas* | COII | *An. melas* | MW179567 |
| SENP2.G2 | *An. gambiae* s.s.*-melas* hybrid | COII | *An. gambiae* s.s. | MW179568 |
| SENP2.A10 | *An. melas* | COII | *An. melas* | MW179569 |
| SENP2.B10 | *An. melas* | COII | *An. melas* | MW179570 |
| SENP2.G10 | *An. melas-gambiae* s.s. hybrid | COII | *An. melas* | MW179571 |
| SENP2.H10 | *An. melas-gambiae* s.s. hybrid | COII | *An. melas* | MW179572 |
| SENP2.G11 | *An. melas* | COII | *An. melas* | MW179573 |
| SENP2.H11 | *An. melas* | COII | *An. melas* | MW179574 |
| SENP2.A12 | *An. melas* | COII | *An. melas* | MW179575 |
| SENP2.B12 | *An. melas* | COII | *An. melas* | MW179576 |
| SENP2.G12 | *An. melas-gambiae* s.s. hybrid | COII | *An. melas* | MW179577 |
| SENP1.F10 | *An. melas* | COII | *An. melas* | MW179578 |
| SENP1.E11 | *An. coluzzii - melas* hybrid | COII | *An. coluzzii* | MW179579 |
| SENP1.H11 | *An. melas-gambiae* s.s. hybrid | COII | *An. melas* | MW179580 |
| SENP2.G1 | *An. melas-gambiae* s.s. hybrid | COII | *An. melas* | MW179581 |
| SENP2.F2 | *An. melas* | COII | *An. melas* | MW179582 |
| FANP2.H2 | *An. gambiae* s.s.*-melas* hybrid | COII | *An. gambiae* s.s. | MW179583 |
| FANP2.F4 | *An. gambiae* s.s.*-melas* hybrid | COII | *An. gambiae* s.s. | MW179584 |
| FANP2.G4 | *An. gambiae* s.s.*-melas* hybrid | COII | *An. gambiae* s.s. | MW179585 |
| MAFP1.C8 | *An. gambiae* s.s. | COII | *An. gambiae* s.s. | MW179586 |
| MAFP1.B9 | *An. coluzzii – gambiae* s.s. hybrid | COII | *An. coluzzii* | MW179587 |
