## Supplementary Table 3 for "Evidence for natural hybridisation and novel *Wolbachia* strain superinfections in the *Anopheles gambiae* complex from Guinea"

| **Sample ID** | **Sample host species** | ***Wolbachia* Strain** | **Accession number** |
| --- | --- | --- | --- |
| SENP2.B9 | *An. melas* | *w*Anga-Guinea-A1 | MW179588 |
| SENP2.H9 | *An. melas* | *w*Anga-Guinea-A2 | MW179589 |
