## Supplementary Table 4 for "Evidence for natural hybridisation and novel *Wolbachia* strain superinfections in the *Anopheles gambiae* complex from Guinea"

| ***Wolbachia* Strain** | **Sample ID** | **Sample host species** | ***wsp*** | ***gatB*** | ***coxA*** | ***hcpA*** | ***ftsZ*** | ***fbpA*** |
| --- | --- | --- | --- | --- | --- | --- | --- | --- |
| *w*Anga-Guinea-A1 | SENP2.B9 | *An. melas* | MW179594 | MW179596 | MW179598 | MW179600 | MW179602 | MW179604 |
| *w*AnsX | SENP2.E2 | *An. species X* | MW179595 | MW179597 | MW179599 | MW179601 | MW179603 | MW179605 |
