## Supplementary figures and images for "Evidence for natural hybridisation and novel *Wolbachia* strain superinfections in the *Anopheles gambiae* complex from Guinea"

### Supplementary Figure 1

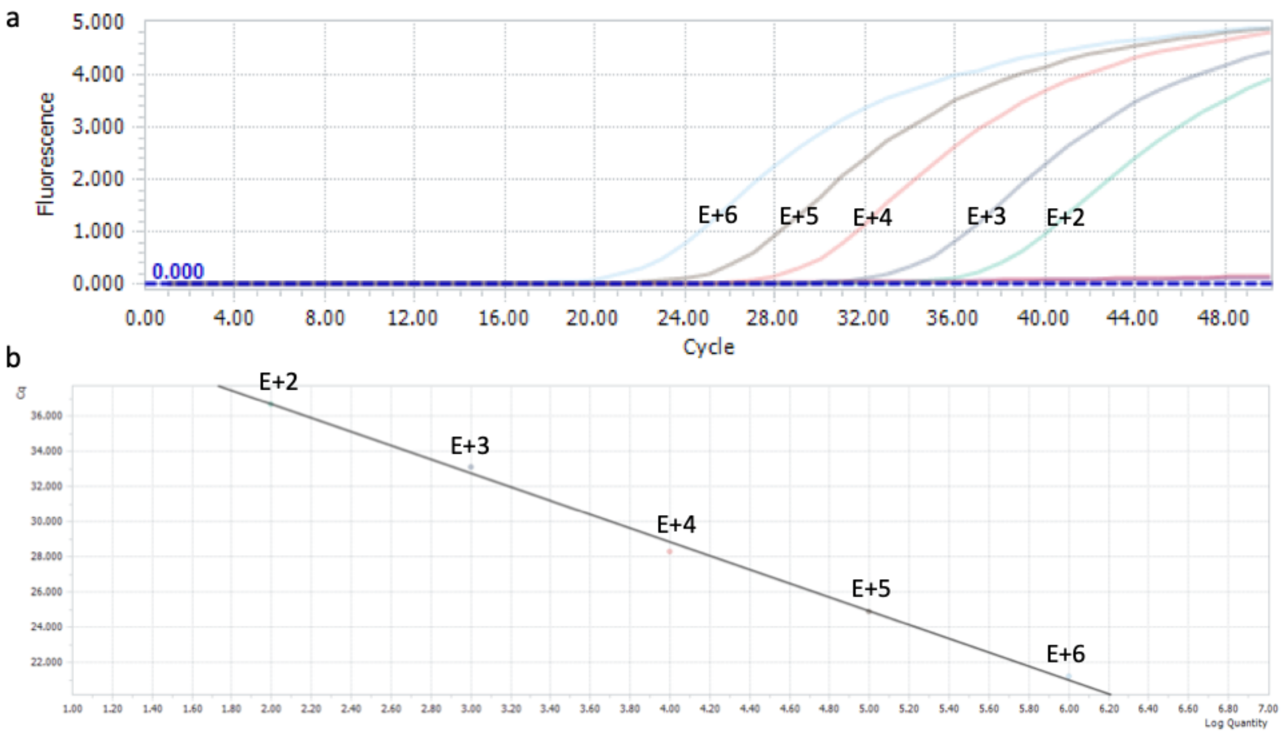
